# A Slow but Steady NanoLuc: R162A mutation results in a decreased, but stable, NanoLuc activity

**DOI:** 10.1101/2023.03.05.531182

**Authors:** Wesam S Ahmed, Anupriya M Geethakumari, Asfia Sultana, Asma Fatima, Angelin M Philip, S. M. Nasir Uddin, Kabir H Biswas

**Affiliations:** Division of Biological and Biomedical Sciences, College of Health & Life Sciences, Hamad Bin Khalifa University, Qatar Foundation, Doha – 34110, Qatar; Division of Genomics and Translational Biomedicine, College of Health & Life Sciences, Hamad Bin Khalifa University, Qatar Foundation, Doha – 34110, Qatar

**Keywords:** Bioluminescence, Luciferase, Molecular Dynamics Simulation, NanoLuc

## Abstract

NanoLuc (NLuc) luciferase has found extensive application in designing a range of biological assays including gene expression analysis, protein-protein interaction and protein conformational changes due to its enhanced brightness and small size. However, questions related to its mechanism of interaction with the substrate, furimazine, as well as bioluminescence activity remains elusive. Here, we combined molecular dynamics (MD) simulation and mutational analysis to show that the R162A mutation results in a decreased but stable bioluminescence activity of NLuc in vitro. Specifically, we performed multiple, all-atom, explicit solvent MD simulations of the apo and furimazine-docked (holo) NLuc structures revealing differential dynamics of the protein in the absence and presence of the ligand. Further, analysis of trajectories for hydrogen bonds (H-bonds) formed between NLuc and furimazine revealed substantial H-bond interaction between R162 and Q32 residues. Mutation of the two residues in NLuc revealed a decreased but stable activity of the R162A, but not Q32A, mutant NLuc in in vitro assays performed with furimazine. In addition to highlighting the role of the R162 residue in NLuc activity, we believe that the mutant NLuc will find wide application in designing in vitro assays requiring extended monitoring of NLuc bioluminescence activity.

## Introduction

Bioluminescence, the phenomenon of light emission during a biochemical reaction, has been effectively taken advantage in the development of a range of biological and biomedical applications. Some of these include gene expression analysis, protein-protein interaction analysis, protein conformational change analysis, blood-based biomarker detection. These applications began with the cloning and utilization of the firefly luciferase (FLuc) in *Escherichia coli*^1^. This was followed by the discovery of the *Renilla* luciferase (RLuc)^2^ as well as *Gaussia* luciferase (GLuc)^3^. While these were useful in developing a variety of bioluminescence-based and Bioluminescence Resonance Energy Transfer (BRET)-based applications such as protein conformational biosensors^4-7^, low bioluminescence activity (brightness) and large sizes of these luciferases led continued research in the area of engineering novel luciferases with better properties. In this regard, structural studies with luciferase derived from deep sea shrimp, *Oplophorus*, revealed a heterotetrametric structure containing two large subunits of 36 kDa and two small subunits of 19 kDa.^2^ Importantly, the discovery of catalytic activity of the smaller subunit led to the idea of using it as a bioluminescent protein independent of the larger subunit.^8, 9^ However, the smaller subunit showed poor expression and stability in cellular environment in the absence of large subunit ^8, 9^. Hall et. al reported a novel luciferase engineered from the small catalytic subunit of *Oplophorus* luciferase with an analogue of luciferase substrate coelenterazine, furimazine^9^. This structurally optimized NanoLuc (NLuc) was small in size (19 kDa) and highly stable at 37°C and showed significant increase in bioluminescence activity compared to the previously reported luciferases such as RLuc and FLuc.^10-13^

Indeed, NLuc has been utilized in a variety of biological and biomedical applications, owing largely to its biophysical properties.^11, 14-17^ Some of these applications include gene regulation analysis^18^, protein aggregation monitoring^19^, protein-protein interaction detection^19, 20^, protein activity monitoring^21^ and cellular imaging^22, 23^. Another area of research where NLuc has been utilized extensively is in the development of protein fragment complementation assay, typically utilized for monitoring protein-protein interaction in living cells.^24-26^ Additional applications of NLuc include nanoparticle conjugates utilized for in vivo imaging^27^, sensitive viral infection detection^28^, to evaluate one of metabolic intermediate in fatty acid metabolism^29^ or to test antibiotic susceptibility^30^. Further, NLuc has been successfully applied in the development of various Bioluminescence Resonance Energy Transfer (BRET)-based biomolecular assays^4-7, 31-33^ such as to study molecular/organelle interactions^34, 35^, for optogenetics purposes^36^, for drug descovery^37, 38^, as genetic encoded light source for a photodynamic anticancer therapy^39^, biomolecule detection^40^, monitoring molecular tension^41^, and assays to monitor SARS-CoV-2 protease activity^42, 43^. Additionally, NLuc possessing synthetic allosteric regulation has been generated through circular permutation of the protein for developing biosensors.^44^ To further expand the utility of NLuc, potent, cell permeable inhibitors have been designed.^45^

While NLuc has been utilized extensively for various applications as listed above, the mechanistic basis for its bioluminescence activity largely remained less understood (Fig. 1A). Structurally, it largely consists of β-sheets (numbered 1 – 11) with four, relatively short α-helices. In the current study, we performed multiple, all-atom, explicit solvent molecular dynamics (MD) simulations with the apo and furimazine-docked structures of NLuc. In addition to revealing differences in the structural dynamics of NLuc in the apo- and holo-forms, analysis of the MD simulation trajectories revealed key hydrogen-bond (H-bond) interactions between R162 and Q32 residues in NLuc with furimazine. We then generated R162A and Q32A mutations in NLuc and performed in vitro assays to determine bioluminescence activity of the proteins revealing a decreased but prolonged bioluminescence of the R162A mutant while similar bioluminescence of the Q32A mutant.

**Fig. 1.**
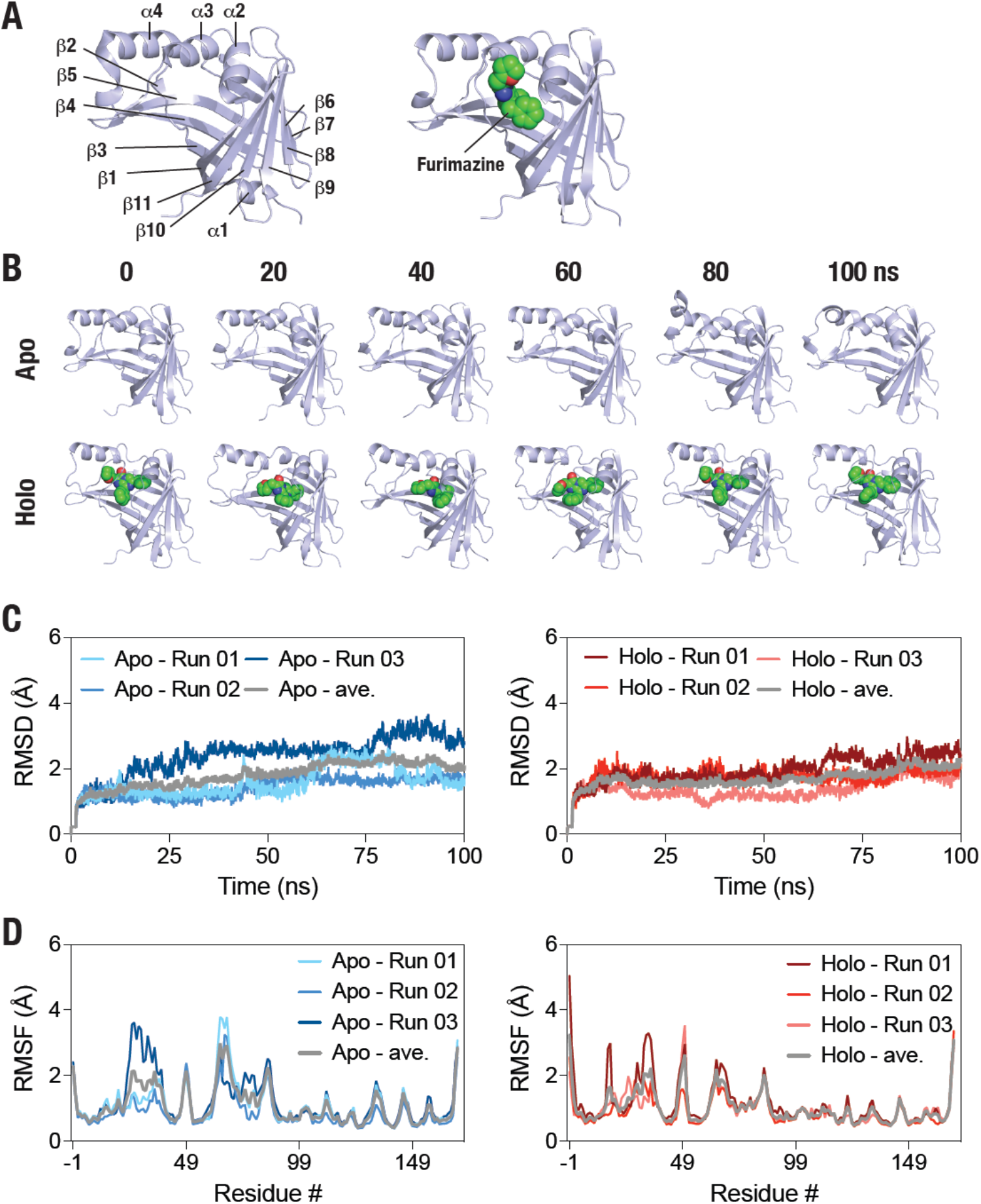
NLuc MD simulations reveals structural stabilization in the presence of substrate furimazine. (A) Cartoon representation of NLuc structure (PBD: 5IB in the apo- (left panel) and holo, furimazine docked (right panel). Secondary structural elements are highlighted in the left panel. (B) Cartoon representation NLuc in the absence (top panel) and in the presence of substrate, furimazin (bottom panel) obtained from a representative 100 ns, all-atom, explicit solvent MD simulation (snapshots acquired every 20 ns). (C-D) Graphs showing Cα RMSD (and RMSF (C) of apo- (left panels) and holo-(right panels) NLuc obtained from 3 independent, 100 ns MD simulations. Grey traces show the average values in each graph.

## Results & Discussions

### *Molecular dynamics simulation reveals dynamic structural* changes in the substrate bound NLuc

In order to understand the structural dynamics of NLuc, we performed all-atom, explicit solvent MD simulations with the apo- and holo, furimazine-bound, NLuc structures (Fig. 1B). For the apo-structure, we used the available crystal structure (PBD: 5IBO) while for the holo-structure, we utilized the furimazine-docked structure of NLuc, which we reported previously^33^. Analysis of the MD simulation trajectories revealed a deeper insertion of the ligand furimazine in the NLuc structure (Fig. 1B). Root-mean-square deviation (RMSD) of Cα atoms of the apo- and holo-NLuc MD simulation trajectories revealed a decrease in the RMSD values of the protein in the presence of the ligand (Fig. 1C) suggesting a furimazine-mediated stabilization of the NLuc structure. Further, root mean square fluctuation (RMSF) analysis of the apo- and holo-NLuc MD simulation trajectories revealed a decrease in fluctuations in residues ranging from 65 to 80 spanning the lid-region of the substrate binding pocket (helix α4) (Fig. 1D) likely indicating furimazine binding-mediated stabilization of the lid-region. Additionally, a decrease in fluctuation was observed for residues ranging from 23 to 30 spanning the loop between helices α2 and α3 as well as residues from 135 to 137 spanning loop between sheets β8 and β9 (Fig. 1D). On the other hand, only residues 18 and 19 located in the beginning of helix α2 showed an increased fluctuation (Fig. 1D).

In order to gain insights into the global structural features of NLuc in the apo- and holo-forms, we performed solvent accessible surface area (SASA) and radius of gyration (RoG) analysis of NLuc over the three MD simulation trajectories (Fig. 2). This revealed reduced SASA in the initial phases (first 50 ns) of the simulation while it reached similar values during the later phase (last 50 ns) of the simulation (Fig. 2A,B). On the other hand, no general alterations were observed in the RoG of the protein in the apo- and holo-forms (Fig. 2C,D). Similarly, no discernable differences were observed in the total, van der Waals and electrostatic energy of the protein in the apo- and holo-forms (Supporting Figure 1). Overall, these analyses revealed a general decrease in the structural dynamics of NLuc in the presence of furimazine.

**Fig. 2.**
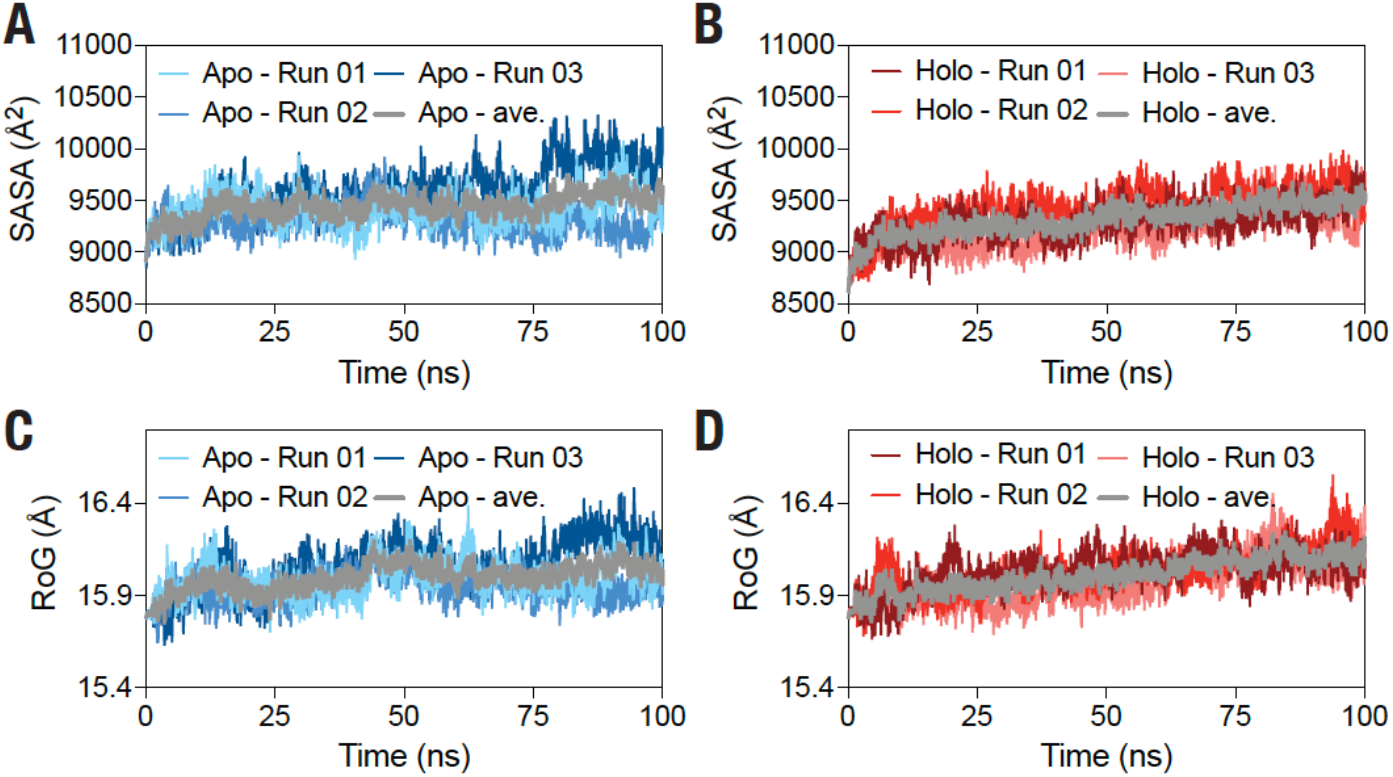
Global structural features of NLuc in the absence and presence of furimazine. Graphs showing SASA (A,B) and RoG (C,D) of the apo (A,C) and holo (B,D) NLuc structures obtained from three independent, 100 ns MD simulations. Grey traces show the average values in each graph.

### Hydrogen bond (H-bond) interactions formed between NLuc and furimazine

We then performed a detailed analysis of the hydrogen bond (H-bond) interactions formed between NLuc and furimazine (Table 1). To limit the analysis to significantly long-lived H-bonds, we applied a 5% cut-off in the fractional occupancy, i.e. life time to select the H-bonds, and determined the mean fractional occupancy of each H-bond interaction. This analysis revealed that the side chain of R162 forms a H-bond with furimazine with the longest fractional occupancy (24.56%) and this H-bond was observed in all three MD simulation runs (Table 1). Second, the sidechain of Q32 also formed H-bond interaction with furimazine with average fractional occupancy of 10.58% and was observed in two out of the three simulation runs (Table 1). All other residues (L18, S28, L92, F31, Q12, G35, V36 and Y109) that formed H-bond interactions, either through their side chains or through main chains, were either observed in one of the three simulation or had shorter fractional occupancy (Table 1).

**Table 1.** H-bonds formed between NLuc and furimazine. bonds formed between NLuc and furimazine with greater than 5% occupancy over each 100 ns MD simulation trajectory are shown in the table.

|  |  | %Occupancy |  |  |  |
| --- | --- | --- | --- | --- | --- |
| # | H-bond | Run 01 | Run 02 | Run 03 | Mean |
| 1 | ARG162-Side-furi1-Side | 22.82 | 12.24 | 38.61 | 24.56 |
| 2 | GLN32-Side-furi1-Side | 0.16 | 20.69 | 10.9 | 10.58 |
| 3 | LEU18-Side-furi1-Side- | 0.2 | 0 | 30.04 | 10.08 |
| 4 | SER28-Side-furi1-Side | 24.32 | 0.04 | 0 | 8.12 |
| 5 | LEU92-Side-furi1-Side | 0.87 | 0.08 | 23.26 | 8.07 |
| 6 | PHE31-Side-furi1-Side | 0.48 | 0.75 | 21.91 | 7.71 |
| 7 | LEU18-Main-furi1-Side | 0.04 | 0 | 15.87 | 5.3 |
| 8 | GLN12-Side-furi1-Side | 6.87 | 3.59 | 0 | 3.49 |
| 9 | GLY35-Main-furi1-Side | 0 | 0.12 | 7.62 | 2.58 |
| 10 | VAL36-Side-furi1-Side | 0 | 0 | 6.36 | 2.12 |
| 11 | TYR109-Side-furi1-Side | 6.2 | 0.04 | 0 | 2.08 |

Given the significant H-bond interactions formed by Q32 and R162, we performed a detailed analysis of the MD simulation trajectories of the holo-NLuc focusing our attention to these residues (Fig. 3A). Measurement of distances between center of mass of NLuc and furimazine in the three MD simulations revealed a stable distance between furimazine and NLuc of ∼10 Å (Fig. 3A) with the distance decreasing in the initial 50 ns of the simulation in one of the runs indicating a deeper insertion of the substrate in NLuc. Distance measurement of Q32 and R162 residues with furimazine also remained constant over the three MD simulation trajectories except for some increase in one of the trajectories (Fig. 3B,C). These results suggest a stable interaction of furimazine with Q32 and R162 residues of NLuc.

**Fig. 3.**
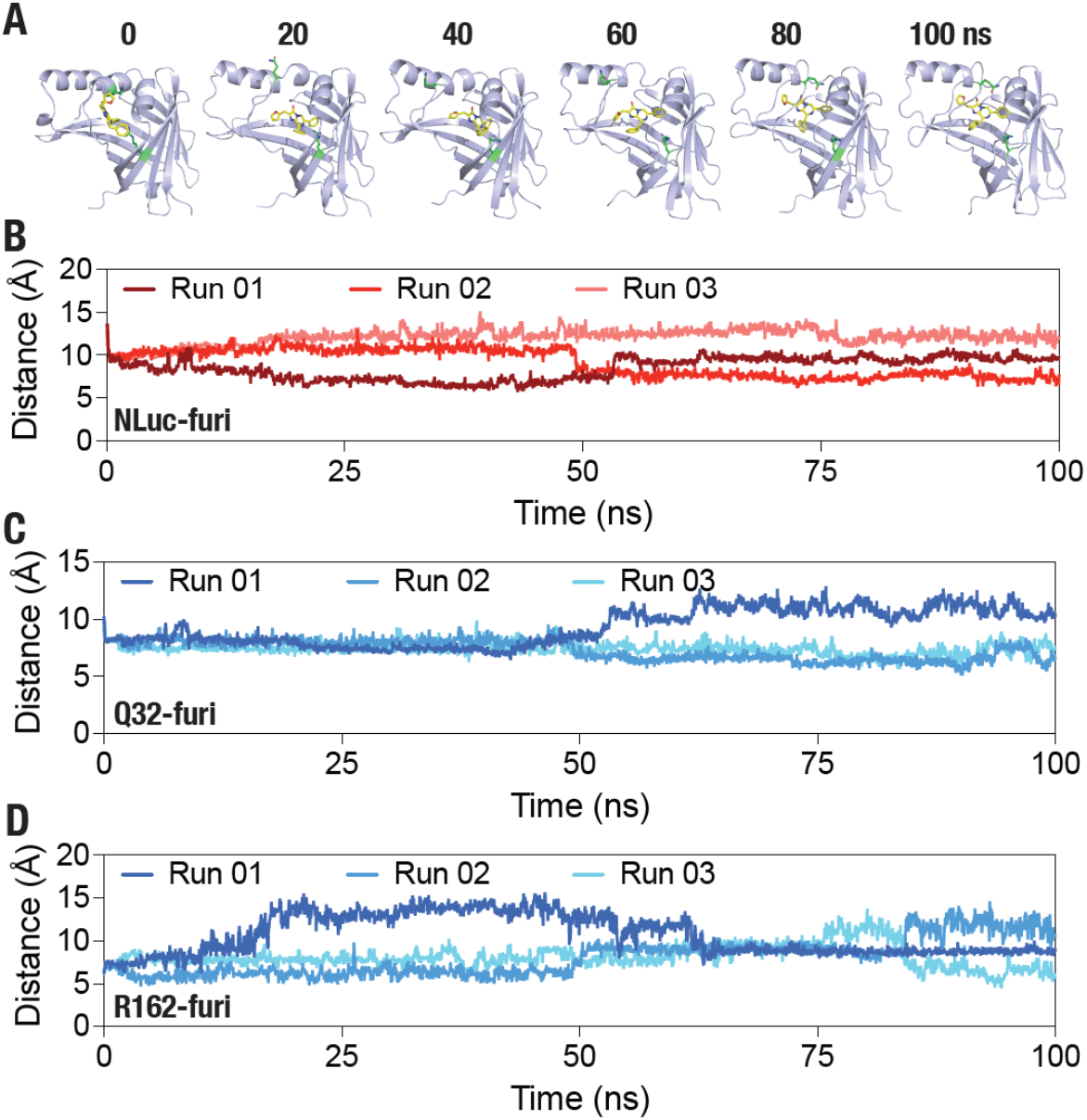
NLuc-furimazine interaction. (A) Cartoon representatio NLuc and furimazine highlighting Q32 and R162 residues that were found to form H-bond interaction with the substrate. (B,C,D) Gra showing center of mass (COM) distance between NLuc furimazine (B), residue 32 and furimazine (C) and R162 a furimazine (D) obtained from three independent 100 ns long, atom, explicit solvent MD simulations.

### R162A mutation in NLuc results in a reduced expression and reduced bioluminescence activity in living cells

Given the stable H-bond interaction formed by residues Q32 and R162 of NLuc with furimazine, we aimed to experimentally determine the impact of mutating these residues into an Ala residue. For this, we generated a fusion protein containing an N-terminal mNeonGreen (mNG) protein and a C-terminal NLuc protein. We expressed the proteins in HEK293T cells and analyzed the protein for expression and bioluminescence activity in living cells. For this, we transfected HEK293T cells and monitored mNG fluorescence to determine expression levels as well as NLuc bioluminescence in the cells over a period of 72 h (Fig. 4). Cells expressing all three constructs showed a time-dependent increase in mNG fluorescence (Fig. 5A). However, the mNG fluorescence was found to be lower in the case of R162A mutant NLuc (Fig. 4A) suggesting a reduced expression of the protein. On the other hand, bioluminescence in cells expressing the WT, Q32A and R162A mutant NLuc constructs did not show such time-dependent increases as seen with mNG fluorescence (Fig. 4B), although lower bioluminescence values were observed in the case of the R162A mutant. These results likely indicate a slower maturation of mNG in the mNG-NLuc constructs as compared to NLuc. Bioluminescence spectra appeared to be similar for all three proteins measured using lysates prepared from transfected cells (Fig. 4C).^32^ Importantly, continuous measurement of NLuc bioluminescence in cells expressing the WT, Q32A and R162A mutant NLuc constructs over a period of 3 h revealed a high but rapidly declining values in the case of WT and Q32A mutant NLuc whereas a reduced but stable bioluminescence was observed in the case of R162A mutant NLuc (Fig. 4D). Indeed, fitting of initial bioluminescence values to an exponential decay model revealed a large increase in the half-life (t_1/2_) of the R162A mutant NLuc compared to the WT and Q32A mutant NLuc (Fig. 4D). In addition to bioluminescence, we monitored BRET (ratio of emission at 533 nm and 467 nm) in cells expressing the WT, Q32A and R162A mutant NLuc and generally found stable except for a larger variation in the case of R162A mutant after 2 h (Fig. 4E).

**Fig. 4.**
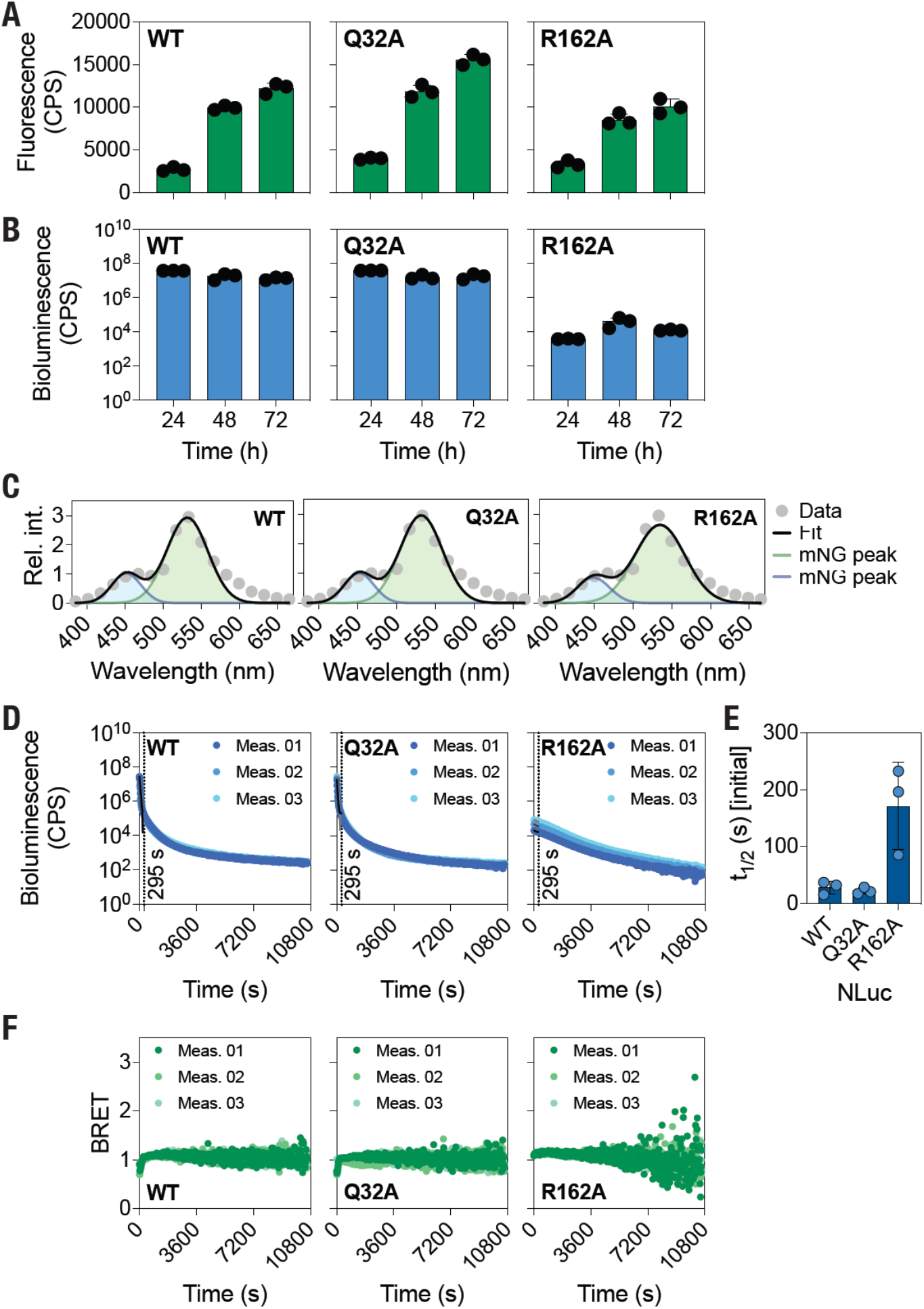
R162A mutation in NLuc results in reduced, but steady, bioluminescence activity. (A,B) Top panel: Schematic representation showing mNG fused NLuc constructs highlighting the Q32A and R162A mutations in NLuc. Bottom panel: Graphs showing fluorescence (A) and bioluminescence (B) values in live cells expressing the WT, Q32A and R162A mutant NLuc fused to mNG after 24, 48 and 72 h of transfection. Data shown are mean ±s.d of three independent measurements, with each measurement performed in duplicate. (C) Graphs showing bioluminescence spectra obtained from lysates prepared from cells expressing WT, Q32A and R162A mutant NLuc fused to mNG. Data were fit to a two-Gaussian model to determine peak NLuc and mNG emission wavelengths. Data shown are mean ± s.d of five measurements. (D) Graphs showing bioluminescence over time in live cells expressing the WT, Q32A & R162A mutant NLuc. Dotted line in each graph at time = 295 s. Data shown are mean ± s.d from three independent measurements, with each measurement performed in duplicates. (E) Graph showing initial bioluminescence activity half-lives of WT, Q32A & R162A mutant NLuc. Data were fit an exponential decay function for the first 295 s (highlighted with a dashed line in the graph). Data shown are mean ± s.d of three independent measurements performed in duplicates. (F) Graphs showing BRET ratio (533 nm over 467 nm) over time in living cells expressing WT, Q32A & R162A mutant NLuc. Data shown are mean ± s.d from three independent measurements, with each measurement performed in duplicates.

**Fig. 5.**
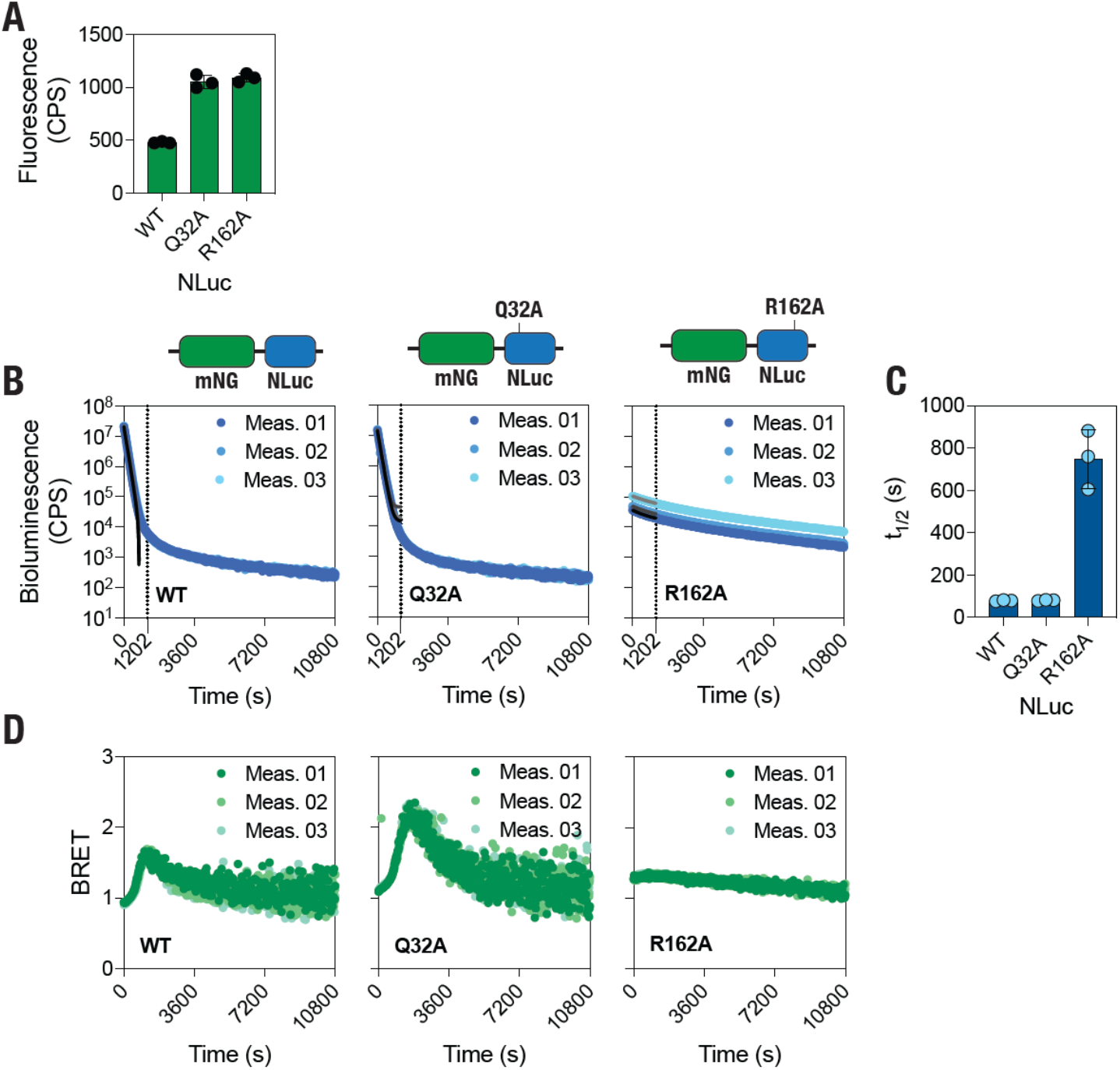
R162A mutation in NLuc results in reduced, but stable, bioluminescence activity in vitro. (A) Graphs showing fluorescence in lysa prepared from cells expressing WT, Q32A & R162A mutant NLuc. Data sho are mean ± s.d of three independent measurements performed in triplicates. Graphs showing bioluminescence over time in lysates prepared from c expressing WT, Q32A & R162A mutant NLuc in vitro. Data shown mean ± s.d from three independent measurements, with each measurement performed in triplicates. (C) Graph showing initial bioluminescence activity half-lives of Q32A & R162A mutant NLuc. Data were fit an exponential decay function for first 1202 s (highlighted with a dashed line in the graph). Data shown are m ± s.d of three independent measurements performed in triplicates. (D) Gra showing BRET ratio (533 nm over 467 nm) over time in lysates prepared fr cells expressing WT, Q32A & R162A mutant NLuc in vitro. Data shown are mean ± s.d obtained from three independent measurements, with each measurem performed in triplicates.

### R162A mutant NLuc shows reduced but stable bioluminescence activity in vitro

Following live cell characterization in terms of bioluminescence and BRET, we characterized the WT and the Q32A and R162A mutant NLuc in vitro. For this, we expressed the mNG-NLuc fusion proteins in HEK293T cells and prepared lysates from the cells after 48 h of transfection (Fig. 5). Measurement of mNG fluorescence in the lysates revealed higher values in the Q32A and R162A mutant NLuc compared to the WT NLuc (Fig. 5A). Continuous monitoring of bioluminescence activity in the lysates revealed a high but a rapidly declining bioluminescence in the case WT and Q32A mutant NLuc while a reduced but stable bioluminescence was observed for the R162A mutant NLuc (Fig. 5B). This is reflected in a much higher initial bioluminescence half-life values obtained from an exponential fitting of the initial (till 1202 s) bioluminescence values of the R162A mutant NLuc compared to the WT and Q32A mutant NLuc (Fig. 5C). These are in agreement with those obtained from live cell experiments. However, BRET values were found to be most stable in the case R162A mutant NLuc compared to the WT and Q32A mutant NLuc (Fig. 5D).

Overall, it appears that R162A mutation in NLuc results in a decrease in the bioluminescence activity of the protein. This can potentially reflect a difference in the interaction of the R162A mutant NLuc with furimazine as H-bond analysis of MD simulation trajectories of the holo-NLuc revealed a relatively stable interaction between R162 and furimazine. In order to test if this is case, we performed three additional all-atom, explicit solvent MD simulations with the holo-R162A mutant NLuc. Determination of free energy changes (ΔG) values of the WT and R162A mutant for furimazine, however, revealed similar values (Fig. 6A). We, therefore, analyzed the trajectories for correlated motions with the proteins by determining dynamic cross-correlation values (Fig. 6B). This revealed regions in the protein that show increased as well as reduced correlation values. For instance, residues ranging from 14 to 42 encompassing parts of sheet β1, helices α2 and α3 and sheet β2 showed increased correlation (Fig. 6B). Similarly, residues ranging from 72 to 84 spanning parts of helix α4 and sheet β4 showed increased correlation. On the other hand, residues in the region from 55 to 68 spanning sheet β3 and the beginning of helix α4 showed reduced correlations. Similarly, residues in the region from 106 and 118 spanning sheet β6 and β7 showed reduced correlations. We note that a structural comparison with another mutant at the same position, R162Q [PDB: 7MJB], with the WT NLuc structure did not reveal any difference in the structure of the protein (Supporting Figure 2). Overall, these results suggest that the R162A mutation does not impact furimazine binding to NLuc. Instead, it affects the bioluminescence activity through a change in the structural dynamics of the protein.

**Fig. 6.**
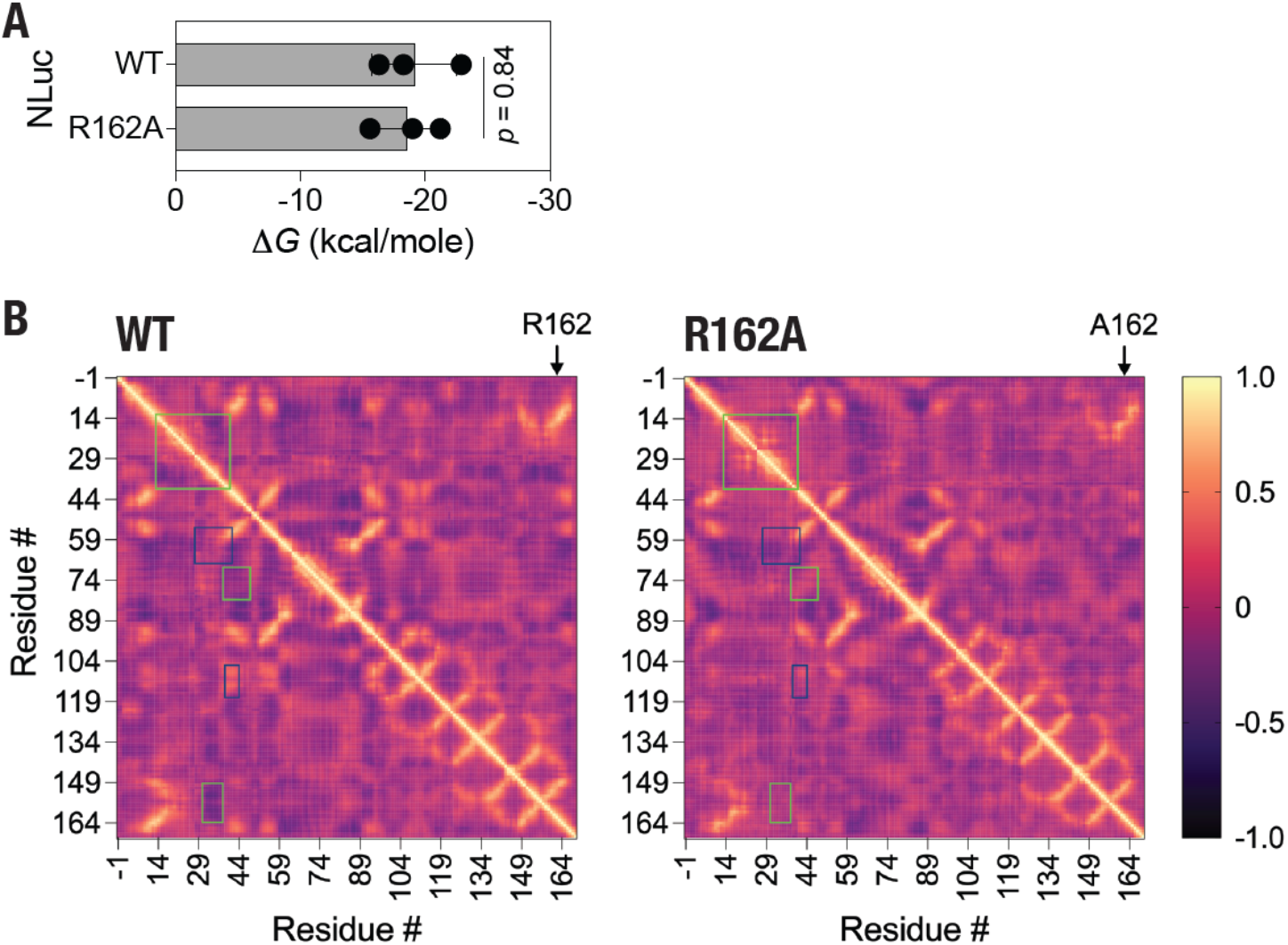
R162A mutation alter NLuc structural dynamics. (A) Graph showing free energy change (ΔG) values of WT and R162A mutant NLuc with furimazi Data shown are mean ± s.d obtained from three independent 100 ns long atom, explicit solvent MD simulations. p value, non-parametric Student’s t-test. (B) Heat maps showing average DCC values of WT (left panel) and R162A mutant (right panel) NLuc obtained from three independent, 100 ns simulations. Positions of residue R162 and A162 are highlighted in each pa Green boxes, increased DCC in the R162A mutant; blue boxes, decreased DCC in the R162A mutant.

We note that while this manuscript was under preparation, a preprint article describing the first furimazine-bound crystal structure of NLuc appeared in the literature^46^, wherein furimazine was shown to bind both to the catalytic site as well as a previously unknown allosteric site. Importantly, authors suggested an involvement of the R162 residue in the protonation of the substrate required for catalytic activity of the enzyme. However, based on the results presented in the current manuscript, it appears that while R162 may serve in the protonation of the substrate, other residues, such as Q12, present in the vicinity could substitute in the R162A mutant through additional rearrangement of residues and substrate in the catalytic site. In a way, it appears that while NLuc is thermally stable, implying enhanced structural rigidity, it is ‘flexible’ catalytically.

## Conclusion

To conclude, we have performed MD simulations of the apo- and holo-NLuc structure docked with the substrate furimazine and showed a general reduction in the structural dynamics of protein in the furimazine bound state. H-bond analysis of the furimazine-bound NLuc structure revealed significant interaction of the substrate with residues R162 and Q32. Mutational analysis of NLuc, wherein the key H-bond forming residues Q32 and R162 were mutated to an Ala revealed a large reduction in the bioluminescence activity of the R162A mutant while no discernable impact was observed for the Q32A mutant. Additional MD simulations of the R162A mutant NLuc and comparison with the WT NLuc revealed no significant changes in the binding interaction of the R162A mutant NLuc with furimazine. However, several differences in the correlated motions in the protein were observed upon R162A mutation. We believe that the results presented here shed light on the structural dynamics and bioluminescence activity of NLuc. Further, given the slow bioluminescence activity of the R162A mutant NLuc leading to prolonged half-life of the bioluminescence activity, the R162A mutant NLuc could be utilized in assays requiring continuous monitoring of bioluminescence activity over long periods without requiring any specialized reagent such as in monitoring biochemical activity of enzymes, including proteases, that display a slow turn-over rate.^31, 32^

## Methods

### NLuc structure preparation and MD simulations

Structural and dynamic effects of furimazine binding to NLuc were investigated using Molecular dynamics (MD) simulation^32, 33, 47-49^ utilizing NAMD 2.13 software^50^ and CHARMM36 force field^51^. Topology and parameter files for apo and holo simulation systems were generated using the CHARMM-GUI online server^52^. For the apo structure, we utilized the crystalized structure of NLuc (PDB: 5IBO, aa: -1 – 169). For the furimazine-bound (holo) structure, we utilized the furimazine-docked NLuc complex that we reported previously^33^ in which the docking was performed on the 5IBO 3D structure. The R162A mutation was modeled on the docked structure using the “Mutagenesis” tool in Pymol (Molecular Graphics System, Version 2.0.0, Schrödinger, LLC; pymol. org).The biomolecular simulation systems were generated by dissolving the 3D structural models in explicit solvent using TIP3P cubic water box^53^ with 10 Å minimum distance between edge of box and any of the atoms in the model. Charges were neutralized by adding 0.15 M NaCl to the solvated system. The biomolecular simulation systems consisted of 78986, 78995, 78982 atoms for apo, holo(WT) and holo(R162A) NLuc, respectively.

Before the production run, the systems underwent energy minimization and thermal equilibration with periodic boundary conditions applied as described previously^47, 54-56^. After equilibration, 100 ns production runs were conducted with a 2 fs timestep, and trajectory frames saved every 10,000 steps. Short-range interactions were handled using a 12 Å cutoff with 10 Å switching distance while long-range electrostatic interactions were computed using the Particle-mesh method with a 1 Å PME grid spacing.

The MD trajectories were analyzed using the tools available in VMD^57^. The backbone RMSD was calculated using the “RMSD-trajectory tool” in VMD, and the RMSF was determined for the Cα atoms of the residues. VMD was also used to calculate the radius of gyration, solvent accessible surface area, and the center-of-mass distance between the paired selections. Hydrogen bond analysis was performed with a cut-off distance of 3.5 Å and an A-D-H angle of 20° using the “Hydrogen Bonds” analysis plugin in VMD. Energy calculations were performed using “NAMD energy” plugin in VMD. The CaFE1.0 analysis tool^58^ was used to estimate the binding free energy changes through the MM-PBSA method^59^. Dynamic cross-correlation (DCC) analysis was based on the position of Cα atoms, and was performed using the Bio3D R package^60 61^.

### NLuc plasmid construct generation

For monitoring NLuc bioluminescence activity and expression levels, we generated a plasmid construct containing an N-terminal mNG, a linker consisting of 3x repeat of the helical peptide EAAK (EAAK)3^62-64^, and a C-terminal NLuc (pmNG-(EAAK)3-NLuc) (see Supporting Text for full nucleotide and amino acid sequences). A gene fragment consisting of parts of mNG, the linker and NLuc was synthesized with BstXI and XhoI sites at the 5’ and 3’ sites respectively (Integrated DNA Technologies, IDT; Iowa, USA). The gene fragment was excised using *Bst*XI and *Xho*I restriction enzymes and then subcloned into similarly digested pmNeonGreen-DEVD-NLuc (Addgene: 98287)^65^ plasmid DNA. NLuc Q32A and R162A mutant plasmid constructs were generated in a similar plasmid construct (GenScript Biotech (Singapore) Pte. Ltd., Singapore). A gene fragment consisting of the NLuc gene was then excised from the above plasmid using *Not*I and *Xho*I restriction enzymes and subcloned into the pmNG-(EAAK)3-NLuc plasmid to generate the pmNG-(EAAK)3-NLuc(Q32A) and pmNG-(EAAK)3-NLuc(R162A) plasmid constructs (see Supporting Text for full nucleotide and amino acid sequences).

### Cell culture, transfection, and live cell experiments

Human embryonic kidney (HEK) 293T cells were maintained in Dulbecco’s modified Eagle’s media (DMEM) with 10% fetal bovine serum (FBS), 1X penicillin-streptomycin at 37 °C in a humidified incubator with an atmosphere of 5% CO_2_.^4-7, 66-73^ Cells were transfected using polyethyleneimine (PEI) lipid (Sigma-Aldrich; 408727-100mL) following the manufacturer’s instructions. Briefly, HEK 293T cells were seeded onto 96-well white plates one day prior to transfection. The plasmid DNA (150 ng/ well) was added to the Opti-MEM (Invitrogen; 31985088), followed by the addition of 1.25 μg/well of PEI lipid and incubated at room temperature for 30 min before being added to cells.

For live cell experiments, HEK293T cells were transfected with pmNG-(EAAK)3-NLuc, pmNG- (EAAK)3-NLuc(Q32A) and pmNG-(EAAK)3-NLuc(R162A) plasmid DNA. After 24 or 48 h of transfection, fluorescence, bioluminescence, and BRET^4-7, 31-33^ measurements were performed by the addition of furimazine (Promega, Wisconsin, USA) at a dilution of 1:200 using a Tecan SPARK® multimode plate reader at 37 °C. For fluorescence measurements, untransfected cells were used to determine blank values, which were subtracted from the fluorescence values of cells expressing the mNG fluorescence proteins. Three independent experiments were performed in triplicates.

### Cell lysate preparation and in vitro assays

HEK 293T cells were seeded onto three 6 cm dishes before 24 h of transfection. The cells were transfected with either NLuc wild-type plasmid (pmNG-(EAAK)3-NLuc) or with NLuc mutant plasmids (pmNG-(EAAK)3-NLuc(Q32A) or pmNG-(EAAK)3-NLuc (R162A)) using polyethylenimine (PEI) lipid. The cell lysate was prepared as described earlier.^32, 74^ Briefly, post 48 h of transfection, the cells were washed with chilled Dulbecco’s Phosphate-Buffered Saline (DPBS) and lysed in a buffer containing 50 mM HEPES (pH 7.5), 50 mM NaCl, 0.1% Triton-X 100, 1 mM dithiothreitol (DTT) & 1 mM ethylenediamine tetraacetic acid (EDTA) on ice, followed by centrifugation at 4 °C for 1 h at 14,000 rotations per min (RPM). The supernatant was collected and stored at -80 °C until further usage. Equal amount of cell lysate, as ascertained from the mNG fluorescence measurements acquired at 530 nm, were used for bioluminescence measurements.^4-7, 32, 33^ After the addition of furimazine at a dilution of 1:200, the bioluminescence (467 nm) and fluorescence (533 nm) readings were recorded at 37 °C using Tecan SPARK multimode microplate reader. BRET^4-7, 31-33^ was determined as a ratio of emissions at 533 nm and 467 nm. For fluorescence measurements, values obtained from wells containing cell lysis buffer only were subtracted from wells containing mNG fluorescent protein-containing cell lysates. Bioluminescence spectra were acquired by measuring emissions at wavelengths ranging from 385 and 665 nm after addition of the furimazine substrate to the lysates at a dilution of 1:200.

### Data Analysis and Figure Preparation

GraphPad Prism (version 8 for MacOS, GraphPad Software, La Jolla California USA; www.graphpad.com), in combination with Microsoft Excel were used for data analysis and graph preparation. Adobe Illustrator was used for assembling all figures.

## Acknowledgements

This work is supported by an internal funding from the College of Health & Life Sciences, Hamad Bin Khalifa University, a member of the Qatar Foundation. Some of the computational research work reported in the manuscript were performed using high-performance computer resources and services provided by the Research Computing group in Texas A&M University at Qatar. Research Computing is funded by the Qatar Foundation for Education, Science and Community Development (http://www.qf.org.qa).

## Author Contributions

K.H.B. conceived experiments. K.H.B, W.S.A., A.M.G., A.S. and A.F. performed the experiments, analyzed data, prepared figure panels, wrote, and approved the manuscript.

## Competing interests

Authors declare no competing interests.

## Notes

### Competing Interest Statement

The authors have declared no competing interest.

