## Supporting Text for "A Slow but Steady NanoLuc: R162A mutation results in a decreased, but stable, NanoLuc activity"

*pmNG-(EAAK)3-NLuc – WT NLuc construct*

Nucleotide sequence

ATGGGAAGTTCACATCATCATCATCATCACTCATCAGGACTGGTGCCACGGGGGTCTCACATGGTCTCCAAGGGGGAAGAGGACAACATGGCCTCTC
TGCCTGCCACACACGAGCTGCACATCTTCGGCAGCATCAATGGCGTGGACTTTGATATGGTGGGCCAGGGCACAGGCAACCCAAATGACGGCTACGA
GGAGCTGAACCTGAAGAGCACCAAGGGCGATCTGCAGTTCTCCCCTTGGATCCTGGTGCCACACATCGGCTATGGCTTTCACCAGTATCTGCCCTAC
CCTGACGGCATGTCCCCCTTCCAGGCCGCCATGGTGGATGGCTCTGGCTACCAGGTGCACAGGACCATGCAGTTTGAGGACGGCGCCTCCCTGACAG
TGAATTACCGGTATACCTACGAGGGCTCTCACATCAAGGGCGAGGCCCAGGTGAAGGGCACAGGCTTCCCAGCCGATGGCCCCGTGATGACAAACTC
TCTGACCGCCGCCGACTGGTGCCGGAGCAAGAAGACCTACCCTAATGATAAGACAATCATCTCCACCTTTAAGTGGTCTTATACCACAGGCAACGGC
AAGCGGTACAGAAGCACAGCCAGAACCACATATACCTTTGCCAAGCCAATGGCCGCCAACTATCTGAAGAATCAGCCCATGTACGTGTTCAGGAAGA
CAGAGCTGAAGCACTCCAAGACCGAGCTGAATTTCAAGGAGTGGCAGAAGGCCTTTACAGACGTGATGGGCATGGATGAGCTGTACAAGGGTACCGA
AGCAGCAGCAAAAGAAGCAGCAGCAAAAGAAGCAGCAGCAAAAGCGGCCGCgATGGTGTTCACACTGGAGGACTTTGTGGGCGATTGGCGGCAGACC
GCCGGCTATAATCTGGATCAGGTGCTGGAGCAGGGCGGCGTGAGCTCCCTGTTCCAGAACCTGGGCGTGAGCGTGACCCCTATCCAGCGGATCGTGC
TGTCCGGCGAGAACGGCCTGAAGATCGACATCCACGTGATCATCCCATACGAGGGCCTGTCTGGCGATCAGATGGGCCAGATCGAGAAGATCTTCAA
GGTGGTGTACCCCGTGGACGATCACCACTTCAAAGTGATCCTGCACTATGGCACCCTGGTCATCGACGGCGTGACCCCCAATATGATCGATTATTTC
GGCCGGCCTTACGAGGGCATCGCCGTGTTTGACGGCAAGAAGATCACCGTGACAGGCACCCTGTGGAACGGCAATAAGATCATCGACGAGAGACTGA
TCAACCCCGATGGCTCCCTGCTGTTCAGGGTGACTATTAACGGAGTGACTGGCTGGCGGCTGTGCGAAAGGATTCTGGCTTGA

Amino acid sequence

MGSSHHHHHHSSGLVPRGSHMVSKGEEDNMASLPATHELHIFGSINGVDFDMVGQGTGNPNDGYEELNLKSTKGDLQFSPWILVPHIGYGFHQYLPY
PDGMSPFQAAMVDGSGYQVHRTMQFEDGASLTVNYRYTYEGSHIKGEAQVKGTGFPADGPVMTNSLTAADWCRSKKTYPNDKTIISTFKWSYTTGNG
KRYRSTARTTYTFAKPMAANYLKNQPMYVFRKTELKHSKTELNFKEWQKAFTDVMGMDELYKGTEAAAKEAAAKEAAAKAAAMVFTLEDFVGDWRQT
AGYNLDQVLEQGGVSSLFQNLGVSVTPIQRIVLSGENGLKIDIHVIIPYEGLSGDQMGQIEKIFKVVYPVDDHHFKVILHYGTLVIDGVTPNMIDYF
GRPYEGIAVFDGKKITVTGTLWNGNKIIDERLINPDGSLLFRVTINGVTGWRLCERILA*

*pmNG-(EAAK)3-NLuc(Q32A) – Q32A NLuc construct*

Nucleotide sequence

ATGGGAAGTTCACATCATCATCATCATCACTCATCAGGACTGGTGCCACGGGGGTCTCACATGGTCTCCAAGGGGGAAGAGGACAACATGGCCTCTC
TGCCTGCCACACACGAGCTGCACATCTTCGGCAGCATCAATGGCGTGGACTTTGATATGGTGGGCCAGGGCACAGGCAACCCAAATGACGGCTACGA
GGAGCTGAACCTGAAGAGCACCAAGGGCGATCTGCAGTTCTCCCCTTGGATCCTGGTGCCACACATCGGCTATGGCTTTCACCAGTATCTGCCCTAC
CCTGACGGCATGTCCCCCTTCCAGGCCGCCATGGTGGATGGCTCTGGCTACCAGGTGCACAGGACCATGCAGTTTGAGGACGGCGCCTCCCTGACAG
TGAATTACCGGTATACCTACGAGGGCTCTCACATCAAGGGCGAGGCCCAGGTGAAGGGCACAGGCTTCCCAGCCGATGGCCCCGTGATGACAAACTC
TCTGACCGCCGCCGACTGGTGCCGGAGCAAGAAGACCTACCCTAATGATAAGACAATCATCTCCACCTTTAAGTGGTCTTATACCACAGGCAACGGC
AAGCGGTACAGAAGCACAGCCAGAACCACATATACCTTTGCCAAGCCAATGGCCGCCAACTATCTGAAGAATCAGCCCATGTACGTGTTCAGGAAGA
CAGAGCTGAAGCACTCCAAGACCGAGCTGAATTTCAAGGAGTGGCAGAAGGCCTTTACAGACGTGATGGGCATGGATGAGCTGTACAAGGGTACCGA
AGCAGCAGCAAAAGAAGCAGCAGCAAAAGAAGCAGCAGCAAAAGCGGCCGCgATGGTGTTCACACTGGAGGACTTTGTGGGCGATTGGCGGCAGACC
GCCGGCTATAATCTGGATCAGGTGCTGGAGCAGGGCGGCGTGAGCTCCCTGTTC**GCG**AACCTGGGCGTGAGCGTGACCCCTATCCAGCGGATCGTGC
TGTCCGGCGAGAACGGCCTGAAGATCGACATCCACGTGATCATCCCATACGAGGGCCTGTCTGGCGATCAGATGGGCCAGATCGAGAAGATCTTCAA
GGTGGTGTACCCCGTGGACGATCACCACTTCAAAGTGATCCTGCACTATGGCACCCTGGTCATCGACGGCGTGACCCCCAATATGATCGATTATTTC
GGCCGGCCTTACGAGGGCATCGCCGTGTTTGACGGCAAGAAGATCACCGTGACAGGCACCCTGTGGAACGGCAATAAGATCATCGACGAGAGACTGA
TCAACCCCGATGGCTCCCTGCTGTTCAGGGTGACTATTAACGGAGTGACTGGCTGGCGGCTGTGCGAAAGGATTCTGGCTTGA

Amino acid sequence

MGSSHHHHHHSSGLVPRGSHMVSKGEEDNMASLPATHELHIFGSINGVDFDMVGQGTGNPNDGYEELNLKSTKGDLQFSPWILVPHIGYGFHQYLPY
PDGMSPFQAAMVDGSGYQVHRTMQFEDGASLTVNYRYTYEGSHIKGEAQVKGTGFPADGPVMTNSLTAADWCRSKKTYPNDKTIISTFKWSYTTGNG
KRYRSTARTTYTFAKPMAANYLKNQPMYVFRKTELKHSKTELNFKEWQKAFTDVMGMDELYKGTEAAAKEAAAKEAAAKAAAMVFTLEDFVGDWRQT
AGYNLDQVLEQGGVSSLF**A**NLGVSVTPIQRIVLSGENGLKIDIHVIIPYEGLSGDQMGQIEKIFKVVYPVDDHHFKVILHYGTLVIDGVTPNMIDYF
GRPYEGIAVFDGKKITVTGTLWNGNKIIDERLINPDGSLLFRVTINGVTGWRLCERILA*

*pmNG-(EAAK)3-NLuc(R162A) – R162A NLuc construct*

Nucleotide sequence

ATGGGAAGTTCACATCATCATCATCATCACTCATCAGGACTGGTGCCACGGGGGTCTCACATGGTCTCCAAGGGGGAAGAGGACAACATGGCCTCTC
TGCCTGCCACACACGAGCTGCACATCTTCGGCAGCATCAATGGCGTGGACTTTGATATGGTGGGCCAGGGCACAGGCAACCCAAATGACGGCTACGA
GGAGCTGAACCTGAAGAGCACCAAGGGCGATCTGCAGTTCTCCCCTTGGATCCTGGTGCCACACATCGGCTATGGCTTTCACCAGTATCTGCCCTAC
CCTGACGGCATGTCCCCCTTCCAGGCCGCCATGGTGGATGGCTCTGGCTACCAGGTGCACAGGACCATGCAGTTTGAGGACGGCGCCTCCCTGACAG
TGAATTACCGGTATACCTACGAGGGCTCTCACATCAAGGGCGAGGCCCAGGTGAAGGGCACAGGCTTCCCAGCCGATGGCCCCGTGATGACAAACTC
TCTGACCGCCGCCGACTGGTGCCGGAGCAAGAAGACCTACCCTAATGATAAGACAATCATCTCCACCTTTAAGTGGTCTTATACCACAGGCAACGGC
AAGCGGTACAGAAGCACAGCCAGAACCACATATACCTTTGCCAAGCCAATGGCCGCCAACTATCTGAAGAATCAGCCCATGTACGTGTTCAGGAAGA
CAGAGCTGAAGCACTCCAAGACCGAGCTGAATTTCAAGGAGTGGCAGAAGGCCTTTACAGACGTGATGGGCATGGATGAGCTGTACAAGGGTACCGA
AGCAGCAGCAAAAGAAGCAGCAGCAAAAGAAGCAGCAGCAAAAGCGGCCGCgATGGTGTTCACACTGGAGGACTTTGTGGGCGATTGGCGGCAGACC
GCCGGCTATAATCTGGATCAGGTGCTGGAGCAGGGCGGCGTGAGCTCCCTGTTCCAGAACCTGGGCGTGAGCGTGACCCCTATCCAGCGGATCGTGC
TGTCCGGCGAGAACGGCCTGAAGATCGACATCCACGTGATCATCCCATACGAGGGCCTGTCTGGCGATCAGATGGGCCAGATCGAGAAGATCTTCAA

GGTGGTGTACCCCGTGGACGATCACCACTTCAAAGTGATCCTGCACTATGGCACCCTGGTCATCGACGGCGTGACCCCCAATATGATCGATTATTTC
GGCCGGCCTTACGAGGGCATCGCCGTGTTTGACGGCAAGAAGATCACCGTGACAGGCACCCTGTGGAACGGCAATAAGATCATCGACGAGAGACTGA
TCAACCCCGATGGCTCCCTGCTGTTCAGGGTGACTATTAACGGAGTGACTGGCTGG**GCG**CTGTGCGAAAGGATTCTGGCTTGA

Amino acid sequence
MGSSHHHHHHSSGLVPRGSHMVSKGEEDNMASLPATHELHIFGSINGVDFDMVGQGTGNPNDGYEELNLKSTKGDLQFSPWILVPHIGYGFHQYLPY
PDGMSPFQAAMVDGSGYQVHRTMQFEDGASLTVNYRYTYEGSHIKGEAQVKGTGFPADGPVMTNSLTAADWCRSKKTYPNDKTIISTFKWSYTTGNG
KRYRSTARTTYTFAKPMAANYLKNQPMYVFRKTELKHSKTELNFKEWQKAFTDVMGMDELYKGTEAAAKEAAAKEAAAKAAAMVFTLEDFVGDWRQT
AGYNLDQVLEQGGVSSLFQNLGVSVTPIQRIVLSGENGLKIDIHVIIPYEGLSGDQMGQIEKIFKVVYPVDDHHFKVILHYGTLVIDGVTPNMIDYF
GRPYEGIAVFDGKKITVTGTLWNGNKIIDERLINPDGSLLFRVTINGVTGW**A**LCERILA*

*Key*:
Green – mNG;
Grey – (EAAK)3 - rigid linker;
Blue – NLuc;
Red, Bold – Mutated amino acid residue;

### Supporting Figures:

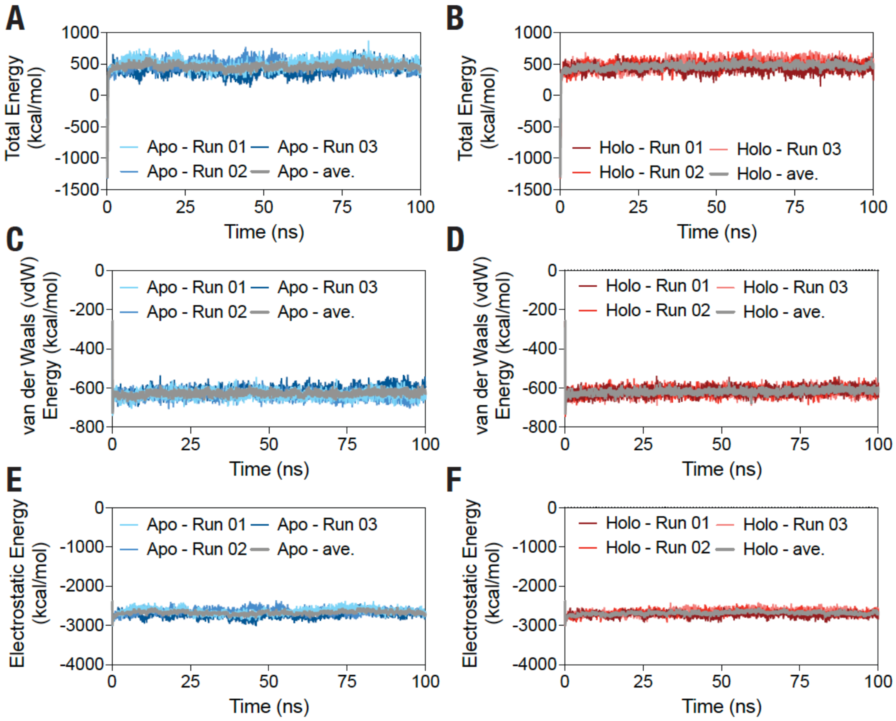


**Supporting Figure 1.** Graphs showing total energy (A,B), van der Waals energy (C,D) and electrostatic energy (E,F) of the apo- (A,C,E) and holo- (B,D,F) NLuc structures obtained from three independent, 100 ns MD simulations. Grey traces show the average values in each graph.

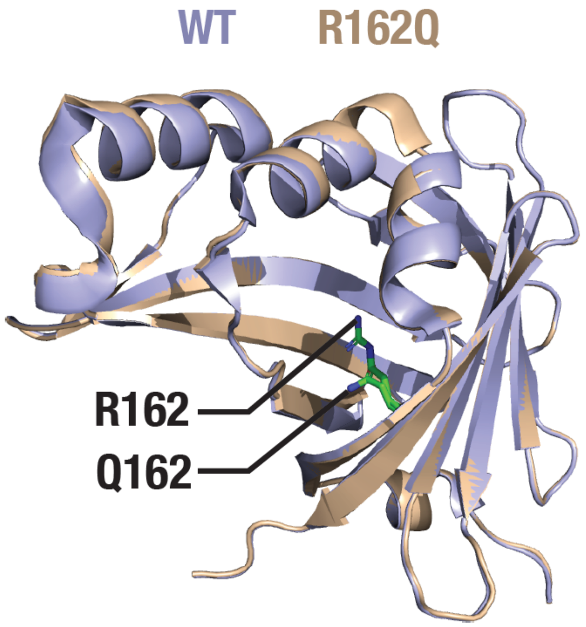


**Supporting Figure 2.** Cartoon representation of aligned WT (PDB: 5IBO) and R162Q mutant (PBD: 7MJB) NLuc structures highlighting the R162 and Q162 residues in stick representation.
